# Caregiving quality modulates neuroendocrine and immunological markers in young children in foster care who have experienced early adversity

**DOI:** 10.1101/2021.04.08.438856

**Authors:** V. Reindl, A. Schippers, K. Tenbrock, A.-K. Job, C. Gerloff, A. Lohaus, N. Heinrichs, K. Konrad

## Abstract

**Background:** Early adversity is believed to alter the body’s stress response systems, putting children at increased risk for somatic and mental health problems. However, it remains unclear whether such alterations normalize under improved caregiving experiences. Thus, the goal of the present study was to investigate i) whether children in foster care show endocrine and immunological alterations relative to children living with their biological families, ii) whether these alterations change over time spent with the foster family and iii) whether the alterations are modulated by current caregiving experiences.

**Methods:** A total of 94 children in foster care and 157 biological children, aged two to seven years, took part in a longitudinal study with three assessments conducted over a 12-month study period. At the initial assessment, children lived for an average of 18 months with their current foster families. Children’s cortisol, dehydroepiandrosterone (DHEA) and progesterone concentrations and cortisol/DHEA ratios were measured in scalp hair and children’s secretory immunoglobulin A (sIgA) levels in saliva. Caregiving quality was assessed based on caregiver-reports and observational measures of caregiver-child interactions.

**Results:** Children in foster care had lower cortisol/DHEA ratios and higher progesterone concentrations than biological children, while no group differences were found for cortisol, DHEA or sIgA. Time spent with the current foster family did not significantly influence the child’s endocrine or immunological markers. Importantly, caregiving quality modulated cortisol/DHEA ratios and sIgA concentrations: children in foster care of lower caregiving quality had lower cortisol/DHEA ratios than children in foster care of higher caregiving quality and showed decreasing, rather than increasing, sIgA concentrations across the study period.

**Conclusions:** Our results indicate that caregiving quality in the foster family may have an important modulating effect on selected indicators of the child’s stress response and could thereby mitigate the possible consequences of early childhood adversity.

## 1. Introduction

Children in foster care typically experience extreme forms of early life stress, including emotional maltreatment, neglect, witnessed violence, and physical and/or sexual abuse in addition to separation from the biological family itself (Oswald, Heil, & Goldbeck, 2009). Such early life adversities have been associated with an increased risk for developmental delays, emotional and behavioral problems, mental disorders, and poor physical health (Oswald et al., 2009; Shonkoff, Boyce, & McEwen, 2009), while out-of-home placement in a stable and caring foster/adoptive family is considered to be an important intervention for reducing such risks (van IJzendoorn & Juffer, 2005).

One possible mechanism linking childhood adversities to later physical and mental health problems may be a dysregulation of the body’s stress response systems, such as the hypothalamic-pituitary-adrenocortical (HPA) axis (McCrory, De Brito, & Viding, 2010). In line with this possibility, studies examining children in institutions, foster or adoption care have found evidence for alterations of the HPA axis with many but not all of them reporting low basal cortisol levels (for a review see: Gunnar & Quevedo, 2008). Such a pattern of hypocortisolism is also supported by a meta-analysis showing a small but significant association between maltreatment and blunted wake-up cortisol levels, though only for samples of agency-referred children, who may have experienced more severe forms of maltreatment. However, no associations have been found with the cortisol awakening response and diurnal cortisol slope (samples include agency-referred and self-report samples, adults and children; Bernard, Frost, Bennett, & Lindhiem, 2017). Such null findings may not necessarily indicate that childhood maltreatment has no or little effect on a child’s HPA axis but could also reflect the considerable heterogeneity of study approaches and findings (see also Fogelman & Canli, 2018). Among several other factors, such as the type, timing and chronicity of maltreatment; the child’s age and gender; psychiatric symptoms; sample recruitment and types of maltreatment assessment, the cortisol assessment method adopted may contribute to this heterogeneity. While cortisol is most commonly sampled in saliva, assessments in scalp hair may provide a better estimate of long-term stress because they reflect cumulative hormone secretion over an extended period of time and are less influenced by diurnal and situational factors (e.g., Stalder et al., 2012; Steudte-Schmiedgen, Kirschbaum, Alexander, & Stalder, 2016). Hair cortisol measurements show considerable stability in the general population (e.g., *r* = .30 - .44 over periods of two to three years in young children; Karlén, Frostell, Theodorsson, Faresjö, & Ludvigsson, 2013). However, it is unclear how cortisol levels develop after foster care placement and during subsequent adjustment to the new family and home environment, which marks a major change in a child’s life. Such longitudinal changes in cortisol concentrations may be moderated by the current quality of caregiving in the foster family.

Evidence, both from animal and human studies, suggests that interactions with his or her caregiver can influence a child’s stress response not only negatively but also positively by acting as a “social buffer”, i.e., dampening the child’s cortisol response to stress (Gunnar & Quevedo, 2008). However, if the caregiver is unable to serve as a reliable social buffer, child HPA axis development may be affected (Flannery, Beauchamp, & Fisher, 2017). At the same time, it seems plausible that under improved caregiving conditions following foster care placement, some of the effects of early childhood adversity on the HPA axis may be reduced. While first cross-sectional and intervention studies support the occurrence of such recovery (DePasquale, Raby, Hoye, & Dozier, 2018; Flannery et al., 2017), other studies indicate that alterations are highly persistent (Kumsta et al., 2017). Research examining longitudinal changes in stress hormone levels after placement in foster/adoption care is scarce.

In addition, most studies examining children’s stress response systems have examined cortisol in isolation. However, many neuroendocrine changes may extend beyond cortisol. Thus, in recent years it has been proposed to move away from single stress-related indicators towards neuroendocrine profiles, e.g., to characterize unique profiles of posttraumatic stress disorder (Michopoulos, Norrholm, & Jovanovic, 2015). In this regard, the hormone dehydroepiandrosterone (DHEA) has received the most attention, since cortisol and DHEA are both released by the activation of the HPA axis (∼ 80% of DHEA and 100% of cortisol are produced by the adrenals) and serve interconnected but largely opposing functions (Kamin & Kertes, 2017). Given this antagonistic dynamic, it may be important to consider each of these indicators not only separately but also in their interplay (cortisol/DHEA ratio), possibly representing an indicator of net glucocorticoid activity (Kamin & Kertes, 2017). For instance, lower morning and afternoon DHEA levels and higher morning and afternoon cortisol/DHEA ratios (measured in saliva) were related to more resilient functioning in a large sample of school-aged children with and without maltreatment experiences (Cicchetti & Rogosch, 2007). Although in this study no effects of maltreatment on hormone concentrations were found, evidence from adult samples suggests that basal DHEA levels are heightened in trauma-exposed individuals (for a review mostly based on plasma assessments, see van Zuiden et al., 2017; for a study on DHEA levels in scalp hair, see Schury et al., 2017).

Another potential candidate is progesterone, a precursor of cortisol biosynthesis (Marceau, Ruttle, Shirtcliff, Essex, & Susman, 2015), which may increase in response to stress, likely to downregulate the stress response (Wirth, 2011). Interestingly, evidence from adults is accumulating linking progesterone to the seeking of social bonds (e.g., Brown et al., 2009; Maner, Miller, Schmidt, & Eckel, 2010), an effect that may be beneficial in coping with stress (Wirth, 2011). While no study that we know of has examined progesterone in children in foster/adoption care or in relation to child maltreatment, given its proposed function in promoting social contact, progesterone may increase when seeking new bonds after foster care placement.

Furthermore, given the link between HPA axis activity and the immune system as well as the heightened risk for somatic health conditions in the aftermath of childhood maltreatment (Kamin & Kertes, 2017; Shonkoff et al., 2009), the investigation of inflammatory markers might contribute to our understanding of the complex neurobiological changes involved and their role in health. One important component of the body’s immune response is immunoglobulin A, which in its secreted form (sIgA) in saliva protects the body from infections of the upper respiratory tract and may serve as a biomarker of stress (Tsujita & Morimoto, 1999). Consistent with the immunosuppressive effects of chronic stress, lower caregiving quality has been found to be associated with reduced sIgA levels in children in the general population (Vermeer, van IJzendoorn, Groeneveld, & Granger, 2012), although the effects of childhood maltreatment remain unknown.

To summarize, growing evidence suggests that childhood maltreatment is associated with alterations to the body’s stress response systems, yet most findings are based on cortisol measures alone and very little is known about the potential for recovery when caregiving improves. Thus, the goal of the current study was to examine whether children in foster care show i) alterations in endocrine and immunological indicators relative to children without documented experiences of maltreatment living with their biological parents and ii) if these alterations change with time spent with the foster family and iii) are modulated by current caregiving experiences. To this end, children in foster care were compared to biological children in the same age range with respect to cortisol, DHEA and progesterone concentrations, measured in scalp hair, the cortisol/DHEA ratio and sIgA levels in saliva. Hormone and sIgA concentrations were measured at three measurement time points, approximately six months apart. Based on previous findings linking childhood maltreatment to a hypoactivation of the HPA axis (Bernard et al., 2017; Gunnar & Quevedo, 2008), we expected children in foster care to show lower cortisol, possibly higher DHEA and lower cortisol/DHEA levels than biological children. In high-quality foster care environments, these alterations may decrease over time, while less or no normalization is expected for children under lower quality care. Given the lack of studies examining sIgA and progesterone in children in out-of-home care, no predictions were formulated in this regard.

## 2. Methods

### 2.1 Study design

Data were collected in the context of a large project on the development of children in foster care (see https://www.uni-bremen.de/klips/forschung/abgeschlossene-drittmittelprojekte/grow-treat for a complete list of the measured variables and Job et al. (2020) for more information on the study design). The study involved a longitudinal assessment conducted at three measurement time points (T1, T2 and T3). With few exceptions, data were collected during home visits, mostly in the afternoon between 1 pm and 4 pm. A parent group training, offered to approximately half of the foster families after T1, had no significant effect on any of the primary and secondary outcome variables, including parenting and caregiver-child interactions, and no effects on additional outcome measures, including hair cortisol or sIgA (Job et al., 2020). Therefore, the intervention was not considered in the subsequent analyses.

### 2.2 Participants

Children in foster care were mainly recruited via youth welfare offices in the surrounding areas of three German cities. Youth welfare agencies were asked to address only long-term, non-kinship care foster families with children aged two to seven years, who had lived in their current foster family for no longer than 24 months and had a history of maltreatment and/or neglect. Biological children in the same age range were recruited mostly via postings or parents’ evenings at nursery and elementary schools. For children in foster care, informed consent was provided by the foster parents and the person(s) holding child custody (i.e., a biological parent, the youth welfare office, a legal guardian or a foster parent). For biological children, informed consent was provided by the biological parents. In a few cases, (foster) parents participated with more than one of their (foster) children. The families were reimbursed for their participation. Ethical approval was obtained by the ethics committee of the German Society for Psychology and additionally of the Medical Faculty of the RWTH Aachen for the biological measures.

The complete sample includes 94 children in foster care (foster care group) and 157 biological children (comparison group) who participated at T1. Children in foster care were aged *M* = 45.64 months (*SD* = 18.82), and biological children were aged *M* = 53.18 months (*SD* = 17.46). Approximately 50% of the sample were female (foster care group: 49%, comparison group: 52%). After T1, 6 children in foster care and 9 biological children and after T2, an additional 4 children in foster care and 2 biological children dropped out of the study. The time between T1 and T2 was *M* = 6.45 months (*SD* = 1.59 months) and the time between T2 and T3 was *M* = 6.04 months (*SD* = 1.49 months). At T1, the children lived for an average of *M* = 17.68 months (*SD* = 8.69 months) with their current foster family. The primary caregiver participated in the study with the child, completed the questionnaires and took part in the caregiver-child interaction observation, which was mostly the (foster) mother. Neither subjects with somatic diseases nor current or past medication were excluded from the current study. However, we controlled in linear mixed model analyses for the effects of asthma/neurodermatitis, which are commonly found chronic diseases in this age range (see also supporting information (SI), Table S1). In addition, the number of infections was assessed at each time point and entered into the models (see section 2.4).

The two groups did not differ with respect to gender distribution, family socioeconomic status (SES), or any health-related variables, including the child’s body mass index (BMI), the prevalence of asthma or neurodermatitis, and the number of infections during the study period (SI, Table S1); however, the children in foster care were significantly younger (*t* (249) = 3.219, *p* = .001, *d* = -.42) and scored higher on (foster) parental ratings of psychopathology (*t* (145.23) *=* -4.456, *p* < .001, *d* = .64). Both groups were used to examine the first and second research objectives, while only the foster care group served as the basis for examining objective 3.

Information on missing data is provided in SI, Table S2. The number of children with at least one valid assessment for cortisol was *N* = 230 (foster care group: *N* = 87, comparison group: *N* = 143), for DHEA and progesterone it was *N* = 241 (foster care group: *N* = 90, comparison group: *N* = 151) and for sIgA it was *N* = 244 (foster care group: *N* = 90, comparison group: *N* = 154). Hair steroid data were mainly missing because children or parents refused the cutting of the child’s hair or the child’s hair had an insufficient amount. sIgA data were mainly missing because children refused the buccal swab.

### 2.3 Assessments

#### 2.3.1 Assessments of hair steroids and sIgA

##### Hair steroid assessments

Two to three hair strands were taken scalp-near from the posterior vertex region. Steroid hormone concentrations were determined in the proximal 3 cm long hair segment, which is assumed to reflect the accumulated hormone secretion of the three months prior to sampling, given a hair growth rate of 1 cm per month (for more information see SI, Text S1). Concentrations of cortisol, DHEA and progesterone were determined by liquid chromatography-tandem mass spectrometry (LC-MS/MS) at TU Dresden following the published protocol of Gao et al. (2013), which is considered the gold standard method for hair steroid hormone analysis.

##### sIgA assessments

A minimum of 30 minutes before saliva collection, the children were asked not to eat or drink anything except for water. Aseptic cotton swabs (Salimetrics, USA) were placed underneath the children’s tongues for 60 s to 90 s and frozen until further analysis. slgA concentrations in saliva were measured using the ready-set-go ELISA system for human IgA (Affymetrix: eBioscience, USA) with a sample dilution of 1:7500 at the University Hospital RWTH Aachen (SI, Text S1).

#### 2.3.2 Assessments of caregiver-child interactions and relationship quality

Parental warmth and support were assessed using the respective subscale of the Zurich Brief Questionnaire for the Assessment of Parental Behaviors (ZKE, Züricher Kurzfragebogen zum Erziehungsverhalten; Reitzle, Winkler Metzke, & Steinhausen, 2001) using the caregiver rating on 12 items (Cronbach’s alpha at T1; foster care group: α = .59, comparison group: α = .77). In addition, relationship quality was assessed by a single item in a caregiver-interview: “*How happy do you consider your relationship with your child?”* on a scale of 1 (“very unhappy’’) to 8 (“perfect”). Finally, in foster families, behavioral coding of nurturing and dysfunctional parenting was derived from an observational measure of caregiver-child interactions (Dyadic Parent–Child Interaction Coding System 4th edition, DPICS IV; Eyberg et al., 2013). To this end, 5 min of child-led play and 5 min of parent-led play were videotaped and coded by independent blinded raters with high interrater reliability (*r*_*tt*_ = .71 to .87; Job et al., 2020).

#### 2.3.3 Assessments of potential confounding and influential factors

Several potential confounds that may influence hair steroid and sIgA measures were considered in the subsequent analyses (for more information see SI, Text S2). Specifically, we included the following sociodemographic variables: child age and gender; family SES based on the social class index by Winkler and Stolzenberg (2009) and whether the child was mainly cared for at home or was enrolled in school, kindergarten or any form of day care. Health-related variables considered include the child’s BMI, a (suspected) diagnosis of asthma or neurodermatitis and the child’s number of (mild) infections during the study period (assessed by caregiver-reports). Maltreatment-related characteristics considered for the foster care group include the types of maltreatment experiences reported by the responsible youth welfare offices and the number of placement changes before placement in the current foster family. Finally, the child’s emotional and behavioral problems were assessed by caregiver-reports using *T* scores for the total score of the German versions of the Child Behavior Checklist (CBCL; CBCL 1½–5 for children aged 2 to 4 years, Achenbach & Rescorla, 2000; CBCL 4–18 for children aged 5 to 9 years, Achenbach, 1991).

### 2.4 Statistical analyses

Statistical analyses were conducted in IBM SPSS Statistics 27 and R version 3.6.2 (R Core Team, 2019; for more information see SI, Text S3). Prior to the analyses, cortisol/DHEA ratios were calculated by dividing cortisol values by DHEA values. Cortisol, DHEA, cortisol/DHEA, progesterone and sIgA values were log-transformed, reducing the positive skewness of the distributions. Then, outlying hair steroid and sIgA values (± 3 SDs from the mean) were identified based on the complete dataset (T1 – T3) and excluded (cortisol: *N* = 5, DHEA: *N* = 6, cortisol/DHEA: N = 4, progesterone: *N* = 8, sIgA: *N* = 6).

First, to examine differences between the foster care and comparison groups, five separate linear mixed models were calculated with cortisol, DHEA, cortisol/DHEA, progesterone and sIgA used as dependent variables. Linear mixed models can accommodate missing values; thus, they allow the researcher to take all available data into account even if only one or two measurement time points are available for the subject. The full model included a random intercept for subjects and the fixed effects of measurement time point (‘time’), group (0 = biological children, 1 = children in foster care) and potential covariates: child age (at T1, in months), gender (0 = female, 1 = male), BMI, the average number of infections per month, asthma/neurodermatitis (0 = no, 1 = yes), SES, day care (0 = no day care, 1 = day care/school) and the CBCL total score. Time was coded as a continuous variable, defined as the time elapsed since T1 in months; thus, for T1, this value was always 0. Continuous variables were centered. Non-significant fixed effects of the potential covariates were reduced backwards based on likelihood-ratio tests.

Second, to test for any differential change between the two groups in hair steroid and sIgA concentrations, the time x group interaction was added to the linear mixed models described above. Furthermore, since the children had lived for an average of ∼ 18 months with their current foster families at T1, separate models were calculated for the foster care group with the predictors of time and time with the current foster family (at T1, in months) as well as the identified covariates.

Third, to explore whether hair steroid and sIgA concentrations were modulated by caregiving experiences in the foster family, we conducted a cluster analysis in the foster care group based on the different caregiver-report and observational measures of the caregiver-child interaction and relationship quality (section 2.3.2), after multiple imputation of missing values. Cluster analysis can consider multiple dimensions simultaneously and may thereby describe meaningful subgroups of families (Henry, Tolan, & Gorman-Smith, 2005). Again, models were calculated with the clustering variable and the identified covariates used as predictors. In addition to the main effect of cluster, we tested for interactive effects between time and cluster in separate models. Furthermore, the results were validated by conducting a cluster analysis in the complete sample based on the available caregiver-report measures and by testing interactive effects between group and cluster. For all research questions, a false discovery rate (FDR) - correction of the *p*-values was applied (Benjamini & Hochberg, 1995), correcting for five tests each (given the five steroid/immunological markers) and in case of significant interactions for the number of post-hoc tests. A detailed description of the statistical/cluster analyses is provided in SI, Text S3, Figure S1. Exploratory analyses examining the effects of maltreatment type and placement changes on hair steroids and sIgA can be found in SI, Text S7.

## 3. Results

First, preparatory analyses examining the stabilities of hair steroid and sIgA concentrations across the study period and their intercorrelations were conducted (SI, Text S4 & S5, Tables S4 – S7). Depending on time distance and variables, the stabilities mostly ranged from *r* = .30 to *r* = .65.

### Children in foster care had lower cortisol/DHEA and higher progesterone concentrations

To examine whether the two groups differed in hair steroid and sIgA concentrations, we calculated linear mixed models with group, time and potential covariates used as predictors (Figure 1; SI, Table S8). In the reduced models, the group effect was found to be significant only for cortisol/DHEA (*t* (178.9) = -2.497, *p* = .013, *p*_*adj*_ = .034) and progesterone (*t* (221.6) = 2.723, *p* = .007, *p*_*adj*_ = .034) but not for cortisol (*t* (170.3) = -1.590, *p* = .11), DHEA (*t* (222.1) = 1.250, *p* = .21) or sIgA (*t* (236.7) = - .677, *p* = .50). These results indicate that the children in foster care had lower cortisol/DHEA and higher progesterone concentrations than the biological children.

**Figure 1.**
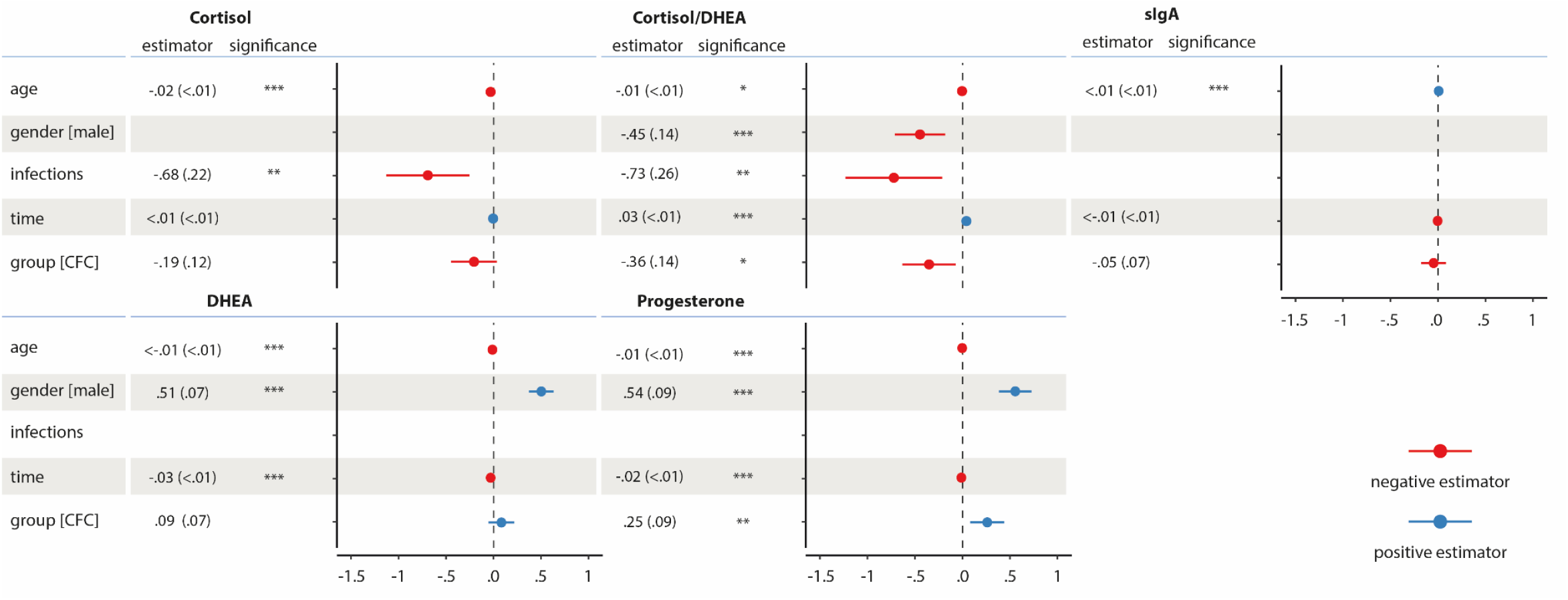
Fixed effect estimates (with standard errors in parentheses) of the linear mixed models predicting cortisol, DHEA, cortisol/DHEA, progesterone and sIgA. Age: child’s age at T1 in months (centered). Gender: 0 = female, 1 = male. Infections: mean number of infections per month across the study period (centered). Time: time elapsed since T1 in months. Group: 0 = biological children, 1 = children in the foster care (CFC). Reduced models are reported after removing non-significant fixed effects of potential covariates. *p < .05, ** p < .01, *** p < .001.

### Time in foster care was not associated with hair steroids or sIgA

Next, we explored whether the two groups displayed differential longitudinal trajectories by adding the time x group interaction to the models (SI, Table S9). Results showed that the time x group interaction did not significantly predict cortisol (*t* (303.8) = 1.075, *p* = .28), DHEA (*t* (406.3) = .755, *p* = .45), progesterone (*t* (359.3) = .585, *p* = .56), cortisol/DHEA (t (307.7) = .853, *p* = .39) and sIgA (*t* (438.3) = .676, *p* = .50). Furthermore, in the foster care group, no significant effect was found for time spent with the current foster family at T1 (cortisol: *t* (57.5) = .795, *p* = .43; DHEA: *t* (80.4) = 1.533, *p* = .13; progesterone: *t* (81.0) = 1.548, *p* = .13; cortisol/DHEA: *t* (58.5) = .702, *p* = .49; sIgA: *t* (84.4) = -.631, *p* = .53) (SI, Table S10).

### Cortisol/DHEA and sIgA concentrations were modulated by caregiving for children in foster care

To identify a measure of caregiving quality that adequately reflects multiple dimensions of parental behavior and caregiver-child relationships and interactions, first, a cluster analysis was conducted in the foster care group based on the different caregiver-report and observational measures. The cluster analysis yielded two subgroups, one with lower caregiving quality and one with higher caregiving quality (SI, Table S11). Groups did not differ on any of the potential covariates (SI, Table S12) except for the CBCL score (*t* (91) = -5.556, *p* <.001, *d* = .79), indicating that children in foster care of lower caregiving quality showed more emotional and behavioral problems than children in foster care of higher caregiving quality.

Examining differences between clusters in hair steroids and sIgA, results showed that children in foster care of lower caregiving quality had lower cortisol levels (*t* (85.86) = -2.173, *p* = .033, *p*_*adj*_ = .081), however, the effect was only marginally significant after FDR-correction, and lower cortisol/DHEA ratios (*t* (61.23) = -2.751, *p* = .008, *p*_*adj*_ = .039) than children in foster care of higher caregiving quality. No effects were found for DHEA, progesterone or sIgA (Figure 2; SI, Table S13). Furthermore, a significant time x cluster interaction was found for sIgA (*t* (92.55) = -2.684, *p* = .008, *p*_*adj*_ = .04) but not for cortisol, DHEA, cortisol/DHEA or progesterone (SI, Table S14). Children in foster care of lower caregiving quality showed decreasing sIgA levels across the study period (*p* = .028, *p*_*adj*_ = .049) while children in foster care of higher caregiving quality showed increasing sIgA levels (*p* = .049, *p*_*adj*_ = .049). Including the CBCL score in the models did not change the main findings.

**Figure 2.**
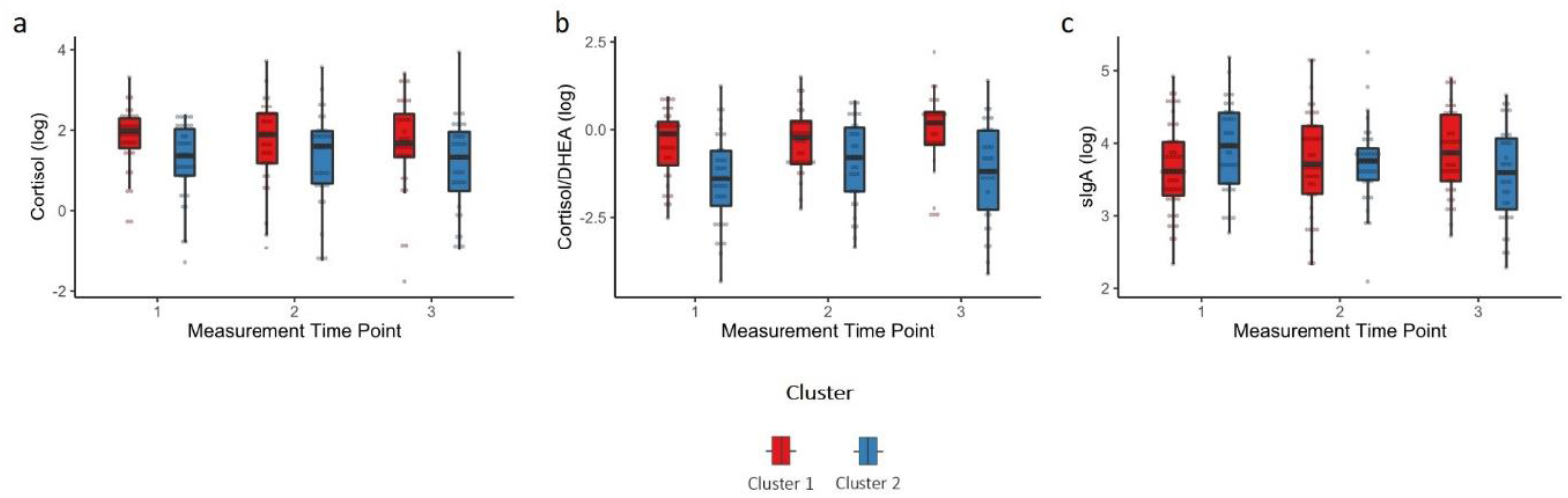
a) Cortisol, b) Cortisol/DHEA and c) sIgA levels (log transformed) of children in foster care of higher caregiving quality (cluster 1) and children in foster care of lower caregiving quality (cluster 2). Boxplots are depicted whereby the median is presented by the horizontal bar, the lower and upper hinges correspond to the 25th and 75th percentiles and the whiskers extend from the hinge to the largest/smallest value no further than the inter-quartile range multiplied by 1.5.

To examine whether these findings were specific to children in foster care, we additionally conducted a cluster analysis in the complete sample based on the caregiver-report measures, which were available from both sample groups (the ZKE subscale and the relationship satisfaction item; SI, Text S6, Tables S15 & S16). Results showed a significant group x cluster interaction for cortisol and cortisol/DHEA: Only in the foster care group was higher caregiving quality found to be associated with higher cortisol and higher cortisol/DHEA concentrations, while no effects of caregiving quality were found in the comparison group. Furthermore, children in foster care of lower caregiving quality had lower cortisol and cortisol/DHEA concentrations than children in biological families of lower caregiving quality, while no group differences were found in the ‘high caregiving quality’ cluster. For sIgA, a marginally significant time x group x cluster interaction emerged, indicating that children in foster care of lower caregiving quality showed decreasing sIgA levels over time, while children in foster care with higher caregiving quality showed sIgA increases. Together, these results support the above findings and indicate that caregiving quality modulated hair steroid and immunological activities only in the foster care group and not in the comparison group.

## 4. Discussion

The current study findings demonstrated lower cortisol/DHEA ratios and higher progesterone levels in children in foster care relative to biological children, while no group differences were observed in terms of cortisol, DHEA or sIgA. Furthermore, time spent with the current foster family at T1 was not found to be associated with hair steroids and sIgA and no significant group differences were found in the longitudinal trajectories. Importantly, our results indicate that children in foster care of lower caregiving quality had marginally lower cortisol concentrations and lower cortisol/DHEA ratios than children in foster care of higher caregiving quality, and showed decreasing, rather than increasing, sIgA concentrations across the study period.

### Effects of early adversity and caregiving on the HPA axis

Lower cortisol concentrations in children in foster care of lower caregiving quality may indicate a hypoactivation of the HPA axis. Our findings, although only marginally significant, are generally in line with theoretical models, suggesting that initially, cortisol levels are elevated after experiencing traumatic events but are eventually attenuated with increasing chronicity, resulting in blunted cortisol levels (Steudte-Schmiedgen et al., 2016). Furthermore, empirically, a significant body of research has reported a hypocortisolism among children in adoption or foster care, mostly using salivary cortisol measures (Bernard et al., 2017; Gunnar & Quevedo, 2008). However, in the current study, the only measure found to be significantly associated with caregiving quality in the foster family was the ratio between cortisol and DHEA, which may thus, represent a more sensitive index of HPA axis activity in children in foster care than either measure alone (see also Kamin & Kertes, 2017). Interestingly, in a study of 6- to 12-year-old children with and without maltreatment experiences, a higher cortisol/DHEA ratio was found to be linked to higher levels of resilient functioning (Cicchetti & Rogosch, 2007).

Importantly, our results indicate that the (marginally) significant associations between caregiving and cortisol and cortisol/DHEA were specific to children in foster care. In line with this finding, a recent meta-analysis showed no significant overall association between parental warmth and children’s morning cortisol levels in observational studies with no reported maltreatment (Hackman, O’Brien, & Zalewski, 2018). Thus, children in foster care could be particularly susceptible to both the positive and negative effects of parenting. Not only could high caregiving quality lead to a normalization of a child’s HPA axis but also could low caregiving quality lead to persistently high levels of stress, resulting in long-term HPA axis hypoactivation.

Furthermore, to date relatively little is known about the role of caregiving on HPA axis activity during adolescence. Preliminary evidence suggests that in general, parental buffering effects decrease during or after puberty when the HPA axis increases in basal activity (e.g., Hostinar, Johnson & Gunnar, 2015), possibly related to pubertal changes. Thus, future studies with longer follow-up assessments of children in foster care after pubertal transition phases are needed to study long-term effects after early adversity.

### Effects of early adversity and caregiving on progesterone

Progesterone has been proposed as a stress-sensitive and stress-responsive hormone that, however, has received very little attention in children, in general, and has so far not been examined in children in foster care. In rodents, the progesterone metabolite allopregnanolone increases social and sexual behaviors (Frye et al., 2006). Furthermore, there is evidence that progesterone shows interrelationships with oxytocin (e.g., Miyamoto & Schams, 1991). Based on these findings, it has been speculated that progesterone, similar to what is posited by the “tend and befriend” hypothesis for oxytocin (Taylor et al., 2000), promotes affiliation as a coping strategy in response to stress (Wirth, 2011). In humans, there is initial evidence to support this, suggesting that progesterone is particularly sensitive to social rejection (in adults; Brown et al., 2009; Maner et al., 2010). Thus, to further clarify the role of progesterone in children in foster care, future studies may examine progesterone levels longitudinally, starting shortly after children are placed in foster care and in relation to the parent’s and child’s developing attachment to each other.

### Relationships to the immune system

Chronic stress and adrenocortical activation are associated with marked alterations in the immune system (Kamin & Kertes, 2017; Segerstrom & Miller, 2004), yet in the current study, we did not find any differences between children in foster care and biological children in terms of sIgA levels, and we found no associations between hair steroids and sIgA (SI, Table S6). One possible explanation is that the effects of current and chronic stress may be powerfully confounded in children in foster care. While sIgA has been shown to increase immediately after stress exposure, it may then gradually decrease (Tsujita & Morimoto, 1999). This delayed stress effect may explain decreasing sIgA levels found in children in foster care of lower caregiving quality across the study period, while children in foster care of higher caregiving quality may show an age-related increase (see also Vermeer et al., 2012).

### Limitations

Some limitations of the study must be acknowledged. First, while the assessment of steroid hormones in hair has several important advantages as a marker of long-term stress, these measurements are relatively novel. The first validation efforts for hair cortisol have yielded promising results, however progesterone and DHEA concentrations in hair must be further validated and explored in child samples (e.g., Gao et al., 2016; Stalder et al., 2012). Second, it is not unlikely that our sample is a selective sample of families who are willing to participate in a longitudinal study with several home visits, resulting in less variability in caregiving quality (see also Job et al., 2020). Third, we cannot draw any causal inferences about the association between early adversity and HPA axis regulation or on the moderating effect of current caregiving. Further, more research is needed to disentangle the effects of caregiving quality and of the associated variable of child behavior problems.

In conclusion, our results contribute to an expanding body of research showing that caregiving experiences, both positive and negative, can shape children’s stress response systems (Gunnar & Quevedo, 2008). Furthermore, our results indicate that the cortisol/DHEA ratio may be a sensitive index of HPA axis functioning in children in foster care, suggesting that future research should focus more on neuroendocrine profiles rather than single indicators.

## Acknowledgments

We are grateful to the families who took part in the study and would like to thank all researchers involved in participant recruitment and data collection, in particular Tabea Symanzik, Sabrina Schütte, Christine Möller and Daniela Ehrenberg.

