## Supporting Information for "Caregiving quality modulates neuroendocrine and immunological markers in young children in foster care who have experienced early adversity"

**Table S1**

Sample description and comparison between children in foster care and biological children in sociodemographic and health-related variables as well as parenting and caregiver-child relationship.

| | *CFC* | *BC* | *Test Statistic* |
|---|---|---|---|
| Sociodemographic Characteristics | | | |
| Female Gender (*n*, %) | 46 (48.9%) | 82 (52.2%) | $X^2$ (1) = .255, *p* = .70 |
| Child's age in months, T1 (*M*, *SD*) | 45.64 (18.82) | 53.18 (17.46) | *t* (249) = 3.219, *p* = .001, *d* = -.42 |
| SES, T1 (*M*, *SD*) | 14.87 (3.73) | 14.95 (4.15) | *t* (242) = .150, *p* = .88 |
| Day care/school, T1 (*n*, %) | 60 (63.8 %) | 154 (98.1%) | $X^2$ (1) = 54.909, *p* < .001, *w* = .47 |
| Physical and Mental Health | | | |
| Neurodermatitis/Asthma (*n*, %) | 17 (18.1%) | 30 (19.1%) | $X^2$ (1) = .040, *p* = .87 |
| Infections per month (*M*, *SD*) | .39 (.31) | .40 (.26) | *t* (201) = .184, *p* = .85 |
| BMI (*M*, S*D*) | 16.14 (1.56) | 16.32 (1.80) | *t* (246) = .797, *p* = .43 |
| CBCL Total Score (*M*, *SD*) | 54.11 (11.65) | 48.01 (8.03) | *t* (145.23) = -4.456, *p* < .001, *d* = .64 |
| Caregiver-Child Interaction and Relationship Quality | | | |
| Relationship Quality (*M*, *SD*) | 6.66 (.98) | 7.02 (.77) | *t* (161.89) = 3.092, *p* = .002, *d* = -.43 |
| Warmth & Support (*M*, *SD*) | 3.66 (.18) | 3.73 (.22) | *t* (245) = 2.291, *p* = .023, *d* = -.30 |
| Nurturing Parenting (*M*, S*D*) | 42.77 (16.77) | | |
| Dysfunctional Parenting (*M*, *SD*) | 13.98 (8.79) | | |

*Note.* CFC: children in foster care; BC: biological children; SES: socioeconomic status. BMI: child body mass index. CBCL: total score of the Child Behavior Checklist (*T*-value). Relationship Quality: assessed by caregiver-interview. Warmth & Support: subscale of the Zurich Brief Questionnaire for the Assessment of Parental Behaviors. Nurturing Parenting and Dysfunctional Parenting: subscales of the Dyadic Parent–Child Interaction Coding System. If not stated otherwise, values were averaged across the three measurement time points.

**Table S2**

Sample sizes for hair steroid and sIgA concentrations of children in foster care and biological children at the three measurement time points

| | | *Cortisol* | *DHEA* | *Progesterone* | *sIgA* | *Cortisol/DHEA* |
|---|---|---|---|---|---|---|
| T1 | *CFC* | 71 | 78 | 74 | 88 | 71 |
| | *BC* | 108 | 111 | 110 | 134 | 107 |
| T2 | *CFC* | 67 | 73 | 73 | 79 | 67 |
| | *BC* | 103 | 116 | 115 | 134 | 103 |
| T3 | *CFC* | 64 | 68 | 68 | 77 | 63 |
| | *BC* | 112 | 123 | 125 | 138 | 107 |

*Note.* CFC: children in foster care; BC: biological children.

**Text S1. Information on hair steroid and sIgA data collection**

**Hair steroid data collection.** All hair strands were stored in aluminium foil and analysed at the end of the study to avoid sampling effects. In most cases, 7.5 mg of whole, non-pulverized hair was used for LC-MS analyses. Samples with less than 4 mg of hair were discarded (12%) while samples with a weight between 4 mg - 7.5 mg were included in the analyses (18.9%). Moreover, a few participants had hair shorter than 3 cm (1.4%). To avoid a reduction of the sample size, these hair strands were included in the analyses. All main results were confirmed in a subsample consisting of hair with 7.5 mg of weight and in a subsample of hair with 3 cm of length.

**sIgA data collection.** Directly after the saliva collection, the swabs were placed in a centrifuge tube, sealed and put in a cooler with icepacks, in which they were transported back to the study site. At the study site, they were frozen at - 20°C (Bielefeld/Braunschweig) or - 80°C (Aachen). Probes were sent with dry ice to the University Hospital RWTH Aachen, where slgA concentrations were analysed in saliva.

**Text S2. Assessments of potential confounding and influential factors**

**Sociodemographic characteristics.** Caregivers provided information on sociodemographic characteristics, including net income, education level and professional position, in a self-developed background interview at T1. Based on this information, the family's socioeconomic status (SES) was calculated based on the social class index by Winkler & Stolzenberg (2009), ranging from 3 to 21 (actual and possible range), with higher vales indicating a higher social class. Based on this index, 16 out of 244 families (6.6%) belonged to the lower social class, 92 families (37.7%) to the middle social class and 136 (55.7%) to the upper social class. Furthermore, caregivers were asked to indicate whether the child was mainly cared for at home or was enrolled in a school, a kindergarten or other forms of day care.

**Health-related variables.** The child's height and weight were measured by research assistants at each of the measurement time points with a standardized weighing scale and measurement tape. The child's body mass index (BMI) was calculated as weight in kg / (height in m)$^2$. Outlying values (> 3 SD of the mean) were removed (T1: $n$ = 3, T2: $n$ = 0, T3: $n$ = 1). Moreover, in a (semi-) standardized interview (Rink, unpublished questionnaire), caregivers reported on the child's health, such as chronic health conditions and the number of infections. The prevalence of asthma and neurodermatitis was summarized to a single score. Asthma / neurodermatitis was assumed to be present if caregivers reported a diagnosed or suspected asthma or neurodermatitis condition at any of the three measurement time points. The number of (mild) infections was reported with respect to the previous three months at T1 and the time since the last assessment at T2 and at T3. Based on this information, the child's average number of infections per month was calculated. If the caregivers reported that the child constantly showed signs of infection, such as a runny nose, this was treated as two episodes per month.

**Experiences of maltreatment and placement-related characteristics.** The number of placement disruptions and the types of maltreatment experiences were reported by the responsible youth welfare offices based on their records (for more information see Ehrenberg, Lohaus, Konrad, & Heinrichs, 2018). Youth welfare officers were asked to state whether the child had experienced sexual abuse, psychological abuse, physical abuse and / or neglect in the family of origin. Information from the youth welfare office was available for 84 children in foster care. Based on the study of Ehrenberg et al. (2018), maltreatment types were grouped into the following categories: no history of maltreatment and neglect, these children were placed in foster care because of inadequate parenting ability ($n$ = 9, 10.7%), experiences of neglect and/or psychological abuse ($n$ = 51, 60.7%) and experiences of neglect, psychological and/or physical/sexual abuse ($n$ = 23, 27.4%). For one child, the type of maltreatment was unknown.

**Child emotional and behavioral problems.** The child's emotional and behavioral problems were assessed by caregiver-report using the German versions of the Child Behavior Checklist (CBCL 1½–5, Achenbach & Rescorla, 2000; CBCL 4–18, Achenbach, 1991). The internal consistency of the total score, measured by Cronbach's alpha, was high for both the foster care group (at T1, CBCL 1½–5: α = .95, CBCL 4–18: α = .95) and the comparison group (at T1, CBCL 1½–5: α = .97; CBCL 4–18: α = .89).

**Text S3. Data analyses**

**Linear mixed models:** Linear mixed model analyses were conducted with the R package 'lme4' (Bates, Mächler, Bolker, & Walker, 2014). Final models were fitted by REML and *p*-values were obtained using the summary function of the R package 'lmerTest' (Kuznetsova, Brockhoff, & Christensen, 2017) with the Satterthwaite approximation for the degrees of freedom.

**Cluster analyses:** First, cluster analyses were conducted for the foster care group based on the ZKE warmth and support subscale, the relationship satisfaction item as well as the two DPICS scales nurturing and dysfunctional parenting at each of the three measurement time points (the procedure is depicted in Figure S1). Second, cluster analyses were repeated for the complete sample (foster care and comparison group) based on the ZKE warmth and support subscale as well as the relationship satisfaction item, again at each measurement time point. The caregiving variables showed mostly high stabilities (except for DPICS dysfunctional parenting; see Table S3).

Prior to the cluster analyses, missing values were imputed using multiple imputation ($N$ = 100) in the R package 'mice' (van Buuren & Groothuis-Oudshoorn, 2011). After the multiple imputation, all variables were standardized by dividing the variables by their ranges (Milligan & Cooper, 1988). Following the suggestions of Basagaña, Barrera-Gómez, Benet, Antó and Garcia-Aymerich (2013), the optimal number of clusters was determined for each imputation. To this end, 30 different indices for determining the number of clusters were calculated using the R package 'NbClust' (Charrad, Ghazzali, Boiteau, & Niknafs, 2014) and the optimal number of clusters was chosen for each imputation according to the majority rule, that is the most frequently found optimal number of clusters was selected. If two number of clusters were found equally often, the lower number of clusters was selected as the optimum. Second, the final number of clusters was chosen as the one appearing with the highest frequency in the imputed data sets, which was two for both cluster analyses. Afterwards, for each of the imputed data sets, for which two was the optimal number of clusters, a k-means cluster analysis was conducted using the R package 'ClusterR' (Mouselimis, 2020) and 'RcppArmadillo', with the 'kmeans++ initialization' (Arthur & Vassilvitskii, 2006), 10 initializations and 300 maximum iterations. Clusters were relabelled so that all have the same meaning. Specifically, a value of 1 was chosen for the cluster with higher caregiving quality and a value of 2 chosen for the cluster with lower caregiving quality. Next, linear mixed models were calculated for each of the imputed data sets. The results of the LMMs were pooled, correcting the degree of freedom according to Barnard and Rubin (1999).

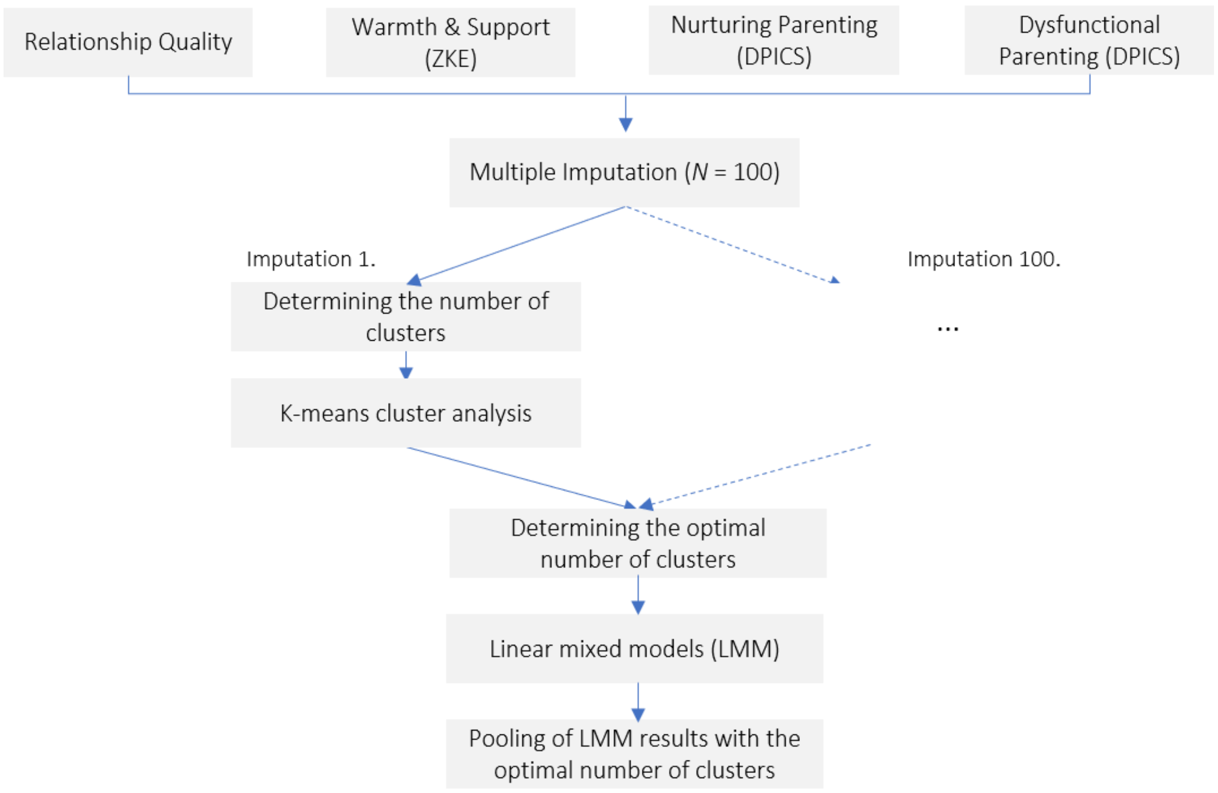


**Figure S1.** Cluster analysis.

**Table S3**

Pearson correlations for caregiving-related study variables between the three measurement time points in the complete sample

| | *T1 – T2* | *T2 – T3* | *T1 – T3* |
|---|---|---|---|
| Relationship Quality | .51 ($p$ < .001) | .63 ($p$ < .001) | .50 ($p$ < .001) |
| Warmth & Support | .50 ($p$ < .001) | .61 ($p$ < .001) | .48 ($p$ < .001) |
| Nurturing Parenting | .60 ($p$ < .001) | .65 ($p$ <.001) | .47 ($p$ < .001) |
| Dysfunctional Parenting | .39 ($p$ = .001) | .46 ($p$ <.001) | .30 ($p$ = .012) |

*Note.* Uncorrected *p*-values are reported in brackets. Relationship Quality: assessed by caregiver-interview. Warmth & Support: subscale of the Zurich Brief Questionnaire for the Assessment of Parental Behaviors. Nurturing Parenting and Dysfunctional Parenting: subscales of the Dyadic Parent–Child Interaction Coding System.

**Text S4. Stabilities of hair steroids and sIgA**

Prior to the main analyses, several preparatory analyses were conducted. First, to examine the stabilities of hair steroid and sIgA concentrations across time, Pearson correlations were calculated between T1 and T2, T1 and T3 as well as T2 and T3 (Table S4). Differences in the correlation coefficients between the foster care and comparison group were examined using Fisher's Z-tests.

In the complete sample, weak to moderate stabilities were found for cortisol ($r$ = .32 – $r$ = .35) and sIgA ($r$ = .41 – $r$ = .53) as well as moderate to strong stabilities for progesterone ($r$ = .59 – $r$ = .63). DHEA concentrations were weakly correlated between T1 and T2 ($r$ = .32, $p$ < .001), moderately correlated between T1 and T3 ($r$ = .58, $p$ < .001) and showed no significant correlation between T2 and T3 ($r$ = .07, $p$ = .40).

**Table S4**

Pearson correlations for child's hair steroid and sIgA concentrations between the three measurement time points in the complete sample

| | *T1 – T2* | *T2 – T3* | *T1 – T3* |
|---|---|---|---|
| Cortisol | .33 ($p$ < .001) | .32 ($p$ < .001) | .35 ($p$ < .001) |
| DHEA | .32 ($p$ < .001) | .07 ($p$ = .40) | .58 ($p$ < .001) |
| Progesterone | .59 ($p$ < .001) | .63 ($p$ < .001) | .59 ($p$ < .001) |
| sIgA | .53 ($p$ < .001) | .52 ($p$ < .001) | .41 ($p$ < .001) |
| Cortisol/DHEA | .32 ($p$ <.001) | .32 ($p$ < .001) | .44 ($p$ < .001) |

*Note.* Uncorrected $p$-values are reported in brackets.

Separate analyses for the two groups can be found in Table S5. Significant differences between the correlation coefficients were found only for progesterone and cortisol/DHEA, as revealed by Fisher's Z-tests. Compared to the comparison group, children in foster care had more stable progesterone concentrations from T1 to T2 ($z$ = 1.876, $p$ = .03) and from T2 to T3 ($z$ = 2.688, $p$ = .004) as well as more stable cortisol/DHEA ratios from T1 to T3 ($z$ = 1.693, $p$ = .045).

**Table S5**

Pearson correlations for child's hair steroid and sIgA concentrations between the three measurement time points in children in foster care and biological children

| | | *T1 – T2* | *T2 – T3* | *T1 – T3* |
|---|---|---|---|---|
| Cortisol | *CFC* | .28 ($p$ = .035) | .32 ($p$ = .024) | .47 ($p$ < .001) |
| | *BC* | .36 ($p$ < .001) | .33 ($p$ = .002) | .26 ($p$ = .017) |
| DHEA | *CFC* | .30 ($p$ = .017) | .01 ($p$ = .92) | .53 ($p$ < .001) |
| | *BC* | .35 ($p$ = .001) | .11 ($p$ = .28) | .60 ($p$ < .001) |
| Progesterone | *CFC* | .68 ($p$ < .001) | .75 ($p$ < .001) | .59 ($p$ < .001) |
| | *BC* | .46 ($p$ < .001) | .47 ($p$ < .001) | .57 ($p$ < .001) |
| sIgA | *CFC* | .51 ($p$ < .001) | .53 ($p$ < .001) | .36 ($p$ = .002) |
| | *BC* | .52 ($p$ < .001) | .49 ($p$ < .001) | .44 ($p$ < .001) |
| Cortisol/DHEA | *CFC* | .17 ($p$ = .21) | .22 ($p$ = .12) | .56 ($p$ <.001) |
| | *BC* | .43 ($p$ <.001) | .41 ($p$ < .001) | .31 ($p$ = .006) |

*Note.* CFC: children in foster care; BC: biological children. Uncorrected *p*-values are reported in brackets.

**Text S5. Associations between hair steroids and sIgA**

To explore the relationships between hair steroids and sIgA concentrations, Pearson correlations were examined. Prior to the analyses, hair steroid and sIgA concentrations were averaged across the three measurement time points. Again, correlation coefficients were compared between the foster care and comparison group using Fisher's Z-tests.

Intercorrelations between the hair steroids and sIgA are shown in Table S6 for the complete sample. Given that hair steroids and sIgA were influenced by child's age and gender, partial correlations were additionally calculated controlling for these confounds ($r_{par}$). Results showed small but significant correlations between cortisol and DHEA ($r_{par}$ = .19, $p$ = .003), moderate correlations between cortisol and progesterone ($r_{par}$ = .34, $p$ < .001) and the highest correlations between DHEA and progesterone ($r_{par}$ = .52, $p$ < .001). No significant correlations were found for sIgA. Separate analyses for the two groups can be found in Table S7. Children in foster care showed a higher correlation between progesterone and cortisol/DHEA than biological children, as indicated by Fisher's *Z*-test ($z$ = 1.843, $p$ = .033).

**Table S6**

Intercorrelations between child's hair steroid and sIgA concentrations across the three measurement time points in the complete sample

| | | *Cortisol* | *DHEA* | *Progesterone* | *sIgA* |
|---|---|---|---|---|---|
| Cortisol | $r$ | - | | | |
| | $r_{par}$ | - | | | |
| DHEA | $r$ | .23 ($p$ < .001) | | | |
| | $r_{par}$ | .19 ($p$ = .003) | | | |
| Progesterone | $r$ | .38 ($p$ < .001) | .64 ($p$ <.001) | | |
| | $r_{par}$ | .34 ($p$ <.001) | .52 ($p$ <.001) | | |
| sIgA | $r$ | -.10 ($p$ = .15) | -.18 ($p$ = .005) | -.23 ($p$ < .001) | |
| | $r_{par}$ | -.02 ($p$ = .76) | -.10 ($p$ = .15) | -.13 ($p$ = .057) | |
| Cortisol/DHEA | $r$ | .77 ($p$ <.001) | -.38 ($p$ < .001) | -.08 ($p$ = .21) | -.02 ($p$ = .82) |
| | $r_{par}$ | .81 ($p$ <.001) | -.33 ($p$ < .001) | -.01 ($p$ = .95) | -.02 ($p$ = .80) |

*Note.* $r$: Pearson correlation. $r_{par}$: partial Pearson correlation, adjusted for child's age and gender. Uncorrected $p$-values are reported in brackets. Prior to the analysis, hair steroid and sIgA concentrations were averaged across measurement time points.

**Table S7**

Partial intercorrelations between child's hair steroid and sIgA concentrations across the three measurement time points in children in foster care and biological children

| | | *Cortisol* | *DHEA* | *Progesterone* | *sIgA* |
|---|---|---|---|---|---|
| Cortisol | *CFC* | - | | | |
| | *BC* | - | | | |
| DHEA | *CFC* | .31 ($p$ = .003) | | | |
| | *BC* | .13 ($p$ = .14) | | | |
| Progesterone | *CFC* | .44 ($p$ < .001) | .44 ($p$ < .001) | | |
| | *BC* | .30 ($p$ < .001) | .58 ($p$ < .001) | | |
| sIgA | *CFC* | .09 ($p$ = .41) | -.02 ($p$ = .85) | -.03 ($p$ = .76) | |
| | *BC* | -.10 ($p$ = .22) | -.12 ($p$ = .14) | -.17 ($p$ = .038) | |
| Cortisol/DHEA | *CFC* | .82 ($p$ < .001) | -.25 ($p$ = .020) | .16 ($p$ = .14) | .08 ($p$ = .50) |
| | *BC* | .81 ($p$ <.001) | -.37 ($p$ < .001) | -.10 ($p$ = .25) | -.09 ($p$ = .30) |

*Note.* CFC: children in foster care; BC: biological children. Partial Pearson correlation coefficients adjusted for child's age and gender are reported. Prior to the analysis, hair steroid and sIgA concentrations were averaged across measurement time points.

**Table S8**

Fixed effect estimates of **time, group** and relevant covariates for linear mixed models predicting hair steroid and sIgA concentrations

| | Cortisol | DHEA | Progesterone | sIgA | Cortisol/DHEA |
|---|---|---|---|---|---|
| Fixed Effect | | | | | |
| Intercept | 1.55 (.09)*** | 1.95 (.06)*** | .21 (.08)** | 3.88 (.05)*** | -.46 (.12)*** |
| Age | -.02 (<.01)*** | <-.01 (<.01)*** | -.01 (<.01)*** | <.01 (<.01)*** | -.01 (<.01)* |
| Gender | - | .51 (.07)*** | .54 (.09)*** | | -.45 (.14)*** |
| Infections | -.68 (.22)** | | | | -.73 (.26)** |
| Time | <.01 (<.01) | -.03 (<.01)*** | -.02 (<.01)*** | < -.01 (<.01) | .03 (<.01)*** |
| Group | -.19 (.12) | .09 (.07) | .25 (.09)** | -.05 (.07) | -.36 (.14)* |
| Random Part | | | | | |
| Intercept | .28 (.53) | .10 (.31) | .32 (.56) | .16 (.41) | .41 (.64) |
| Residual | .74 (.86) | .37 (.61) | .29 (.54) | .20 (.44) | .93 (.96) |

*Note.* For the fixed effects, estimates are presented with the standard errors in parenthesis. For the random part, variances are presented with the standard deviations in parenthesis. Age: child's age at T1 in months (centered). Gender: 0 = female, 1 = male. Infections: mean number of infections per month (centered). Time: time elapsed since T1 in months. Group: 0 = biological children, 1 = children in the foster care. Reduced models are reported after removing non-significant fixed effects of potential covariates. **p* < .05, ** *p* < .01, *** *p* < .001.

**Table S9**

Fixed effect estimates of **time, group**, **time x group** interaction as well as relevant covariates for linear mixed models predicting hair steroid and sIgA concentrations

| | Cortisol | DHEA | Progesterone | sIgA | Cortisol/DHEA |
|---|---|---|---|---|---|
| Fixed Effect | | | | | |
| Intercept | 1.59 (.10)*** | 1.97 (.07)*** | .22 (.08)** | 3.89 (.05)*** | -.42 (.13)** |
| Age | -.02 (<.01)*** | <-.01 (<.01)*** | -.01 (<.01)*** | <.01 (<.01)*** | -.01 (<.01)* |
| Gender | - | .51 (.07)*** | .54 (.09)*** | - | -.45 (.14)*** |
| Infections | -.68 (.22)** | - | - | - | -.73 (.26)** |
| Time | <-.01 (.01) | -.03 (<.01)*** | -.02 (<.01)*** | <-.01 (<.01) | .03 (.01)* |
| Group | -.31 (.16) | .04 (.09) | .21 (.11)* | -.07 (.08) | -.46 (.19)* |
| Time x Group | .02 (.02) | <.01 (<.01) | <.01 (<.01) | <.01 (<.01) | .02 (.02) |
| Random Part | | | | | |
| Intercept | .28 (.53) | .10 (.31) | .32 (.56) | .17 (.41) | .41 (.64) |
| Residual | .74 (.86) | .37 (.61) | .29 (.54) | .20 (.44) | .93 (.96) |

*Note.* For the fixed effects, estimates are presented with the standard errors in parenthesis. For the random part, variances are presented with the standard deviations in parenthesis. Age: child's age at T1 in months (centered). Gender: 0 = female, 1 = male. Infections: mean number of infections per month (centered). Time: time elapsed since T1 in months. Group: 0 = biological children, 1 = children in the foster care. *$p < .05$, ** $p < .01$, *** $p < .001$.

**Table S10**

Fixed effect estimates of **time with foster family** as well as relevant covariates for linear mixed models predicting hair steroid and sIgA concentrations of children in foster care

| | Cortisol | DHEA | Progesterone | sIgA | Cortisol/DHEA |
|---|---|---|---|---|---|
| Fixed Effect | | | | | |
| Intercept | 1.20 (.26)*** | 1.87 (.14)*** | .17 (.19) | 3.85 (.13)*** | -1.03 (.32)** |
| Age | -.02 (<.01)*** | <-.01 (<.01)* | -.02 (<.01)*** | <.01 (<.01) | -.01 (<.01)* |
| Gender | - | .52 (.11)*** | .71 (.15)*** | - | -.42 (.25) |
| Infections | -.47 (.36) | - | - | - | -.38 (.41) |
| Time | .01 (.01) | -.02 (<.01)** | -.02 (<.01)** | <-.01 (<.01) | .04 (.01)** |
| Time in Family | .01 (.01) | <.01 (<.01) | .01 (<.01) | <-.01 (<.01) | .01 (.01) |
| Random Part | | | | | |
| Intercept | .36 (.60) | .08 (.29) | .39 (.63) | .18 (.43) | .43 (.66) |
| Residual | .77 (.88) | .41 (.64) | .28 (.53) | .21 (.46) | 1.04 (1.02) |

*Note.* For the fixed effects, estimates are presented with the standard errors in parenthesis. For the random part, variances are presented with the standard deviations in parenthesis. Age: child's age at T1 in months (centered). Gender: 0 = female, 1 = male. Infections: mean number of infections per month (centered). Time: time elapsed since T1 in months. Time in Family: time spent with current foster family at T1. *$p < .05$, ** $p < .01$, *** $p < .001$.

**Table S11**

Sample description of children in foster care of higher caregiving quality (cluster 1) and children in foster care of lower caregiving quality (cluster 2) with respect to caregiving variables

| | *Cluster 1* (*n* = 53) | *Cluster 2* (*n* = 41) |
|---|---|---|
| Relationship Quality (*M*, *SD*) | | |
| T1 | 7.29 (.72) | 6.03 (1.05) |
| T2 | 7.34 (.67) | 5.72 (.94) |
| T3 | 7.43 (.58) | 5.49 (.78) |
| Warmth & Support (*M*, *SD*) | | |
| T1 | 3.73 (.19) | 3.61 (.25) |
| T2 | 3.68 (.24) | 3.55 (.24) |
| T3 | 3.74 (.20) | 3.62 (.24) |
| Nurturing Parenting (*M*, *SD*) | | |
| T1 | 48.21 (18.37) | 43.28 (16.32) |
| T2 | 45.67 (20.54) | 40.00 (20.12) |
| T3 | 42.48 (20.18) | 33.14 (16.92) |
| Dysfunctional Parenting (*M*, *SD*) | | |
| T1 | 12.95 (7.78) | 15.50 (11.54) |
| T2 | 12.07 (8.47) | 17.65 (14.57) |
| T3 | 10.95 (9.70) | 15.54 (13.30) |

*Note.* Relationship Quality: assessed by caregiver-interview. Warmth & Support: subscale of the Zurich Brief Questionnaire for the Assessment of Parental Behaviors. Nurturing Parenting and Dysfunctional Parenting: subscales of the Dyadic Parent–Child Interaction Coding System. For each child, the most frequently observed cluster assignment across the *N* = 100 multiple imputations was selected for descriptive analyses.

**Table S12**

Sample description of children in foster care of higher caregiving quality (cluster 1) and children in foster care of lower caregiving quality (cluster 2) with respect to potential covariates / influential factors

| | *Cluster 1* (*n* = 53) | *Cluster 2* (*n* = 41) | *Test Statistic* |
|---|---|---|---|
| Sociodemographics | | | |
| Female Gender (*n*, %) | 29 (54.7%) | 17 (41.5%) | $X^2$ (1) = 1.625, *p* = .22 |
| Child's age in months, T1 (*M*, *SD*) | 42.91 (18.25) | 49.17 (19.19) | *t* (92) = -1.614, *p* = .11 |
| Social Class (*M*, *SD*) | 15.26 (3.67) | 14.39 (3.79) | *t* (89) = 1.107, *p* = .27 |
| Day care / school, T1 (*n*, %) | 32 (60.4%) | 28 (68.3%) | $X^2$ (1) = .627, *p* = .52 |
| Physical and mental health | | | |
| Neurodermatitis/Asthma (*n*, %) | 11 (20.8%) | 6 (14.6%) | $X^2$ (1) = .585, *p* = .59 |
| Infections per month (*M*, *SD*) | .40 (.31) | .38 (.31) | *t* (68) = .214, *p* = .83 |
| BMI (*M*, S*D*) | 16.17 (1.39) | 16.11 (1.76) | *t* (91) = .161, *p* = .87 |
| CBCL (*M*, *SD*) | 48.94 (10.16) | 60.68 (10.07) | *t* (91) = -5.556, *p* < .001, *d* = .79 |
| Risk and Protective Factors | | | |
| Time in Foster Family, T1 (*M*, *SD*) | 16.81 (8.23) | 18.80 (9.23) | *t* (92) = -1.105, *p* = .27 |
| Placement Changes (*M*, *SD*) | 1.22 (1.12) | 1.49 (1.46) | *t* (84) = -.939, *p* = .35 |
| Type of Maltreatment (*n*, %) | | | $X^2$ (2) = .324, *p* = .89 |
| No maltreatment experiences | 6 (12.5%) | 3 (8.6 %) | |
| Neglect and/or emotional abuse | 29 (60.4 %) | 22 (62.9 %) | |
| Neglect, emotional and/or physical/sexual abuse | 13 (27.1 %) | 10 (28.6 %) | |

*Note.* SES: socioeconomic status. BMI: child body mass index. CBCL: total score of the Child Behavior Checklist (*T*-value). For each child, the most frequently observed cluster assignment across the *N* = 100 multiple imputations was selected for descriptive analyses. If not stated otherwise, values were averaged across the three assessments.

**Table S13**

Fixed effect estimates of **time, cluster** as well as relevant covariates for linear mixed models predicting hair steroid and sIgA concentrations of children in foster care

| | Cortisol | DHEA | Progesterone | sIgA | Cortisol/DHEA |
|---|---|---|---|---|---|
| Intercept | 1.63 (.17)*** | 1.94 (.11)*** | .47 (.14)*** | 3.78 (.08)*** | -.53 (.24)* |
| Age | -.02 (<.01)*** | <-.01 (<.01)* | -.01 (<.01)** | <.01 (<.01) | -.01 (<.01)* |
| Gender | - | .51 (.11)*** | .77 (.16)*** | - | -.31 (.22) |
| Infections | -.53 (.33) | - | - | - | -.44 (.36) |
| Time | .01 (.01) | -.02 (<.01)** | -.02 (<.01)** | <-.01 (<.01) | .04 (.01)** |
| Cluster | -.51 (.24)* | .23 (.13) | -.17 (.17) | -.02 (.12) | -.80 (.29)** |

*Note.* For the fixed effects, estimates are presented with the standard errors in parenthesis. Linear mixed model results of the multiple imputations were pooled. Age: child's age at T1 in months (centered). Gender: 0 = female, 1 = male. Infections: mean number of infections per month (centered). Time: time elapsed since T1 in months. CBCL: total score of the Child Behavior Checklist, *T*-value. Cluster: 0 = children in foster care of higher caregiving quality, 1 = children in foster care of lower caregiving quality. *$p < .05$, ** $p < .01$, *** $p < .001$.

**Table S14**

Fixed effect estimates of **time, cluster, time x cluster** as well as relevant covariates for linear mixed models predicting hair steroid and sIgA concentrations of children in foster care

| | Cortisol | DHEA | Progesterone | sIgA | Cortisol/DHEA |
|---|---|---|---|---|---|
| Intercept | 1.68 (.20)*** | 1.94 (.12)*** | .47 (.14)** | 3.68 (.09)*** | -.50 (.26) |
| Age | -.02 (<.01)*** | <-.01 (<.01)* | -.01 (<.01)** | <.01 (<.01) | -.01 (<.01)* |
| Gender | - | .51 (.11)*** | .77 (.16)*** | - | -.31 (.22) |
| Infections | -.53 (.33) | - | - | - | -.44 (.36) |
| Time | <.01 (.01) | -.02 (<.01)* | -.02 (<.01)* | .02 (<.01) | .04 (.02) |
| Cluster | -.63 (.31)* | .22 (.17) | -.17 (.19) | .22 (.15) | -.86 (.36)* |
| Time x Cluster | .02 (.03) | <.01 (.02) | <-.01 (.01) | -.04 (.01)** | <.01 (.03) |

*Note.* For the fixed effects, estimates are presented with the standard errors in parenthesis. Linear mixed model results of the multiple imputations were pooled. Age: child’s age at T1 in months (centered). Gender: 0 = female, 1 = male. Infections: mean number of infections per month (centered). Time: time elapsed since T1 in months. CBCL: total score of the Child Behavior Checklist, *T*-value. Cluster: 0 = children in foster care of higher caregiving quality, 1 = children in foster care of lower caregiving quality. *$p$ < .05, ** $p$ < .01, *** $p$ < .001.

**Text S6. Cluster analysis in the complete sample**

To examine whether the associations between hair steroids and caregiving were specific to the foster care group, we additionally conducted a cluster analysis in the complete sample based on the available caregiver-report measures (the ZKE warmth and support subscale and the relationship satisfaction item). Again, two clusters were identified, one with lower and one with higher relationship quality (Table S15). LMM results revealed a significant interaction between group and cluster for cortisol ($t$ (416.87) = -2.463, $p$ = .014, $p_{adj}$ = .035) and cortisol/DHEA ($t$ (394.57) = -2.999, $p$ = .003, $p_{adj}$ = .014) (Table S16). Splitting up the interaction by group, only in the foster care group, a significant effect of cluster was found for cortisol and cortisol/DHEA: children in foster care of lower caregiving quality had lower cortisol ($t$ (131.02) = -1.997, $p$ = .048, $p_{adj}$ = .096) and lower cortisol/DHEA concentrations ($t$ (132.26) = -3.479, $p$ < .001, $p_{adj}$ = .003) than children in foster care of higher caregiving quality. Furthermore, splitting up the interaction by cluster, a group effect was found only for the 'low caregiving quality' cluster: children in foster care of lower caregiving quality had lower cortisol ($t$ (123.63) = -2.388, $p$ = .018, $p_{adj}$ = .074) and lower cortisol/DHEA concentrations ($t$ (116.45) = -2.838, $p$ = .005, $p_{adj}$ = .011) than children in biological families of lower caregiving quality.

In subsequent models, we examined whether the interaction between time, group and cluster predicted hair steroids and sIgA. Only for sIgA, a significant three-way interaction was observed ($t$ (570.99) = -2.349, $p$ = .019, $p_{adj}$ = .095), which was however only marginally significant after FDR-correction. Splitting up the three-way interaction, a highly significant two-way interaction between time and cluster emerged for the foster care group ($t$ (222.76) = -3.465, $p$ < .001, $p_{adj}$ = .002), however not for the comparison group. Within the foster care group, children in cluster 1 (higher caregiving quality) showed increasing sIgA levels across the study period ($t$ (122.84) = 2.511, $p$ = .013, $p_{adj}$ = .023) while children in cluster 2 (lower caregiving quality) showed decreasing sIgA levels ($t$ (99.07) = -2.413, $p$ = .018, $p_{adj}$ = .023).

**Table S15**

Sample description of children with higher caregiving quality (cluster 1) and children with lower caregiving quality (cluster 2) across the foster care and comparison group

| | *Cluster 1* (*n* = 177) | *Cluster 2* (*n* = 74) |
|---|---|---|
| Relationship Quality (*M*, *SD*) | | |
| T1 | 7.33 (.74) | 6.04 (1.07) |
| T2 | 7.24 (.69) | 5.85 (1.05) |
| T3 | 7.30 (.66) | 5.81 (.88) |
| Warmth & Support (*M*, *SD*) | | |
| T1 | 3.78 (.20) | 3.54 (.28) |
| T2 | 3.75 (.19) | 3.47 (.24) |
| T3 | 3.79 (.19) | 3.53 (.27) |

*Note.* Relationship Quality: assessed by caregiver-interview. Warmth & Support: subscale of the Zurich Brief Questionnaire for the Assessment of Parental Behaviors. For each child, the most frequently observed cluster assignment across the *N* = 100 multiple imputations was selected for descriptive analyses.

**Table S16**

Fixed effect estimates of **time, group, cluster, group x cluster** as well as relevant covariates for linear mixed models predicting hair steroid and sIgA concentrations in the complete sample

| | Cortisol | DHEA | Progesterone | sIgA | Cortisol/DHEA |
|---|---|---|---|---|---|
| Intercept | 1.50 (.09)*** | 1.94 (.07)*** | .17 (.08)* | 3.88(.05)*** | -.51 (.12)*** |
| Age | -.02 (<.01)*** | <-.01 (<.01)*** | -.01 (<.01)*** | <.01 (<.01)*** | -.01 (<.01)** |
| Gender | - | .50 (.07)*** | .55 (.09)*** | - | -.42 (.13)** |
| Infections | -.67 (.22)** | - | - | - | -.72 (.25)** |
| Time | <.01 (<.01) | -.03 (<.01)*** | -.02 (<.01)*** | <-.01 (<.01) | .03 (<.01)*** |
| Group | .06 (.16) | <-.01 (.09) | .30 (.12)* | -.05 (.09) | .05 (.18) |
| Cluster | .23 (.19) | .07 (.12) | .18 (.15) | -.04 (.11) | .18 (.22) |
| Group x Cluster | -.66 (.27)* | .13 (.17) | -.21 (.22) | .04 (.15) | -.93 (.31)** |

*Note.* For the fixed effects, estimates are presented with the standard errors in parenthesis. Linear mixed model results of the multiple imputations were pooled. Age: child's age at T1 in months (centered). Gender: 0 = female, 1 = male. Infections: mean number of infections per month (centered). Time: time elapsed since T1 in months. CBCL: total score of the Child Behavior Checklist, *T*-value. Group: 0 = biological children, 1 = children in the foster care. Cluster: 0 = children with higher caregiving quality, 1 = children with lower caregiving quality. *$p < .05$, ** $p < .01$, *** $p < .001$.

**Text S7. Effects of maltreatment type and placement changes**

To examine the effects of maltreatment type and placement changes on hair steroid and sIgA measures, five linear mixed models were fitted, one for each of the outcome variables. Maltreatment type (coded with 0 = no maltreatment experiences, 1 = neglect and/or emotional abuse, 2 = neglect, emotional and/or physical/sexual abuse), number of placement changes (coded as a continuous variable), measurement time point ('time') and relevant covariates were included in the models as predictors. Models were fitted with REML and $p$-values derived from the $F$-statistic, using the 'anova' function and Satterthwaite's approximation (in the 'lmerTest' R package). Results revealed no significant effects of maltreatment type (cortisol: $F$ (2, 48.11) = 1.640, $p$ = .20; DHEA: $F$ (2, 68.81) = .228, $p$ = .80; cortisol/DHEA: $F$ (2, 48.98) = 1.920, $p$ = .16; progesterone: $F$ (2, 72.19) = .240, $p$ = .79; sIgA: $F$ (2, 72.85) = .290, $p$ = .75) and no significant effects of placement changes (cortisol: $F$ (1, 47.76) = .025, $p$ = .88; DHEA: $F$ (1, 74.68) = 1.168, $p$ = .28; cortisol/DHEA: $F$ (1, 49.08) = .183, $p$ = .67; progesterone: $F$ (1, 76.14) = .819, $p$ = .37; sIgA: $F$ (1, 76.07) = .001, $p$ = .97).
